# Impact of temperature on *Legionella pneumophila*, its protozoan host cells, and the microbial diversity of the biofilm community of a pilot cooling tower

**DOI:** 10.1101/2019.12.12.874149

**Authors:** Adriana Torres Paniagua, Kiran Paranjape, Mengqi Hu, Émilie Bédard, Sébastien Faucher

## Abstract

*Legionella pneumophila (Lp)* is a waterborne bacterium known for causing Legionnaires’ Disease, a severe pneumonia. Cooling towers are a major source of outbreaks, since they provide ideal conditions for *Lp* growth and produce aerosols. In such systems, *Lp* typically grow inside protozoan hosts. Several abiotic factors such as water temperature, pipe material and disinfection regime affect the colonization of cooling towers by *Lp.* The local physical and biological factors promoting the growth of *Lp* in water systems and its spatial distribution are not well understood. Therefore, we built a lab-scale cooling tower to study the dynamics of *Lp* colonization in relationship to the resident microbiota and spatial distribution. The pilot was filled with water from an operating cooling tower harboring low levels of *Lp*. It was seeded with *Vermamoeba vermiformis*, a natural host of *Lp*, and then inoculated with *Lp.* After 92 days of operation, the pilot was disassembled, the water was collected, and biofilm was extracted from the pipes. The microbiome was studied using *16S rRNA* and *18S rRNA* genes amplicon sequencing. The communities of the water and of the biofilm were highly dissimilar. The relative abundance of *Legionella* in water samples reached up to 11% whereas abundance in the biofilm was extremely low (≤0.5 %). In contrast, the host cells were mainly present in the biofilm. This suggest that *Lp* grows in host cells associated with biofilm and is then released back into the water following host cell lysis. In addition, water temperature shaped the bacterial and eukaryotic community of the biofilm, indicating that different parts of the systems may have different effects on *Legionella* growth.

## 1. Introduction

*Legionella pneumophila* (*Lp*) is a gram negative, intracellular, waterborne pathogen known for causing Legionnaires disease (LD), a severe pneumonia, contracted by the inhalation of contaminated aerosols (Buse et al., 2012; Fields, 1996; Fliermans, 1996; McDade et al., 1979). *Lp* is the main cause of waterborne disease in the United States with an incident rate of 1.89 cases per 100,000 inhabitants in 2015 (Centers for Disease Control and Prevention, 2018). The estimated annual cost of hospitalization due to LD in the United States exceeds $716 million USD per year (Giambrone, 2013; Whiley et al., 2014). The incidence of outbreaks of LD is on the rise; the CDC reported that between 2000 and 2014, there was an increase of 286% in cases of LD and Pontiac fever in the United States (Centers for Disease Control and Prevention, 2015). A similar trend was reported in Europe (Beauté, 2013; Beauté, 2017).

*Lp* is a natural inhabitant of many aquatic ecosystems such as lakes, hot springs and rivers (Borella et al., 2005; Carvalho et al., 2008; Fliermans et al., 1981; Lin et al., 2007; Ortiz-Roque and Hazen, 1987; Sheehan et al., 2005). There, it can be found as an intracellular parasite of free living amoeba and ciliates (Fields et al., 2002; Rowbotham, 1980). Importantly, *Legionella* is ubiquitous in engineered water systems (Alary and Roy, 1992). *Legionella* has been detected in pools, water fountains, dental units, humidifiers, domestic potable water distribution systems, cooling towers, hospital and hotel hot water systems (Atlas et al., 1995; Hampton et al., 2016; Kyritsi et al., 2018; Leoni et al., 2018; Leoni et al., 2001; Llewellyn et al., 2017; Moran-Gilad et al., 2012; Paranjape et al., 2020; Pereira et al., 2017; Smith et al., 2015; Stout et al., 1992).

The first recognized outbreak of LD that sickened 182 people in 1976 in Philadelphia was associated to a contaminated cooling tower (Kurtz et al., 1982; McDade et al., 1979). Since then, cooling towers have been reported as the source of several outbreaks of LD (Addiss et al., 1989; Bell et al., 1996; Breiman et al., 1990; Brown et al., 1999; Fitzhenry et al., 2017; Greig et al., 2004; Isozumi et al., 2005; Mitchell et al., 1990; Shelton et al., 1994; Wang et al., 2014).

Currently, cooling towers are a major source of outbreaks and cause up to 28% of sporadic cases of LD (Fitzhenry et al., 2017). This is due to the large amounts of aerosols produced by these towers, which are dispersed over long distances of up to 6 km (Beauté, 2017; Bhopal et al., 1991; Cunha et al., 2016; Fisman et al., 2005; Klaucke et al., 1984; Nguyen et al., 2006).

Understanding the conditions affecting growth of *Lp* in water is critical to elucidate the risk factors linked to outbreaks and improve monitoring and management of water systems. *Legionella* spp. can be detected at low levels in the majority of cooling towers; however, promoting factors are required for *Legionella* to reach sanitary risk levels (Llewellyn et al., 2017). Several physical and chemical factors contributing to *Legionella* colonization have been identified. Temperatures between 25 °C and 50 °C are optimal for *Lp* growth and proliferation (Bedard et al., 2015; Katz et al., 2009; Wadowsky et al., 1985; Yamamoto et al., 1992). A long-term study conducted by Pereira et al. (2017), in which the microbiome of the water of a cooling tower was analysed, confirmed that temperature is highly correlated with the presence of *Legionella*. Moreover, the material of the pipes greatly influence the abundance of *Legionella* in water systems and some material, such as PVC, promotes the presence of *Lp* (Buse et al., 2014; Moritz et al., 2010; Proctor et al., 2017; Rogers et al., 1994b; van der Kooij et al., 2005). The use of disinfectant also impacts the presence of *Lp*. In many countries, cooling towers are under surveillance and management plans are carried out to prevent the proliferation of *Legionella* (Kim et al., 2002; McCoy et al., 2012; Springston and Yocavitch, 2017; Whiley, 2016; WHO, 2007).

Biotic factors also affect the presence of *Lp* in cooling towers. High heterotrophic plate counts (HPC) in poorly managed water distribution systems seem to increase the odds of colonization of *Lp* (Messi et al., 2011; Serrano-Suarez et al., 2013). In contrast, some cooling towers that have high HPC do not harbour *Lp*, suggesting that they may host a microbial population resistant to *Legionella* colonization (Duda et al., 2015). The presence of some organisms such as *Cyanobacteria* (Tison et al., 1980) and *Flavobacterium*, (Wadowsky and Yee, 1983) contribute to the growth of *Lp*. Interestingly, other bacteria such as *Pseudomonas* and *Staphylococcus warneri* seem to have an antagonistic effect on the proliferation of *Legionella* (Guerrieri et al., 2008; Hechard et al., 2006; Paranjape et al., 2020). Therefore, the growth and proliferation of *Legionella pneumophila* in water systems seem to be impacted by the resident microbes. The identity and relative abundance of these microbes is influenced by several parameters. The microbial population residing in cooling towers is shaped by local climate and water sources (Llewellyn et al., 2017; Paranjape et al., 2020). Additionally, the microbiota is affected by the disinfectant residuals and application schedule (Hwang et al., 2012; Paranjape et al., 2020). An important limitation of these studies is that they focus on the microbiota of the water. Biofilm plays a crucial role in *Legionella* proliferation and survival (Cooper and Hanlon, 2010; Flemming et al., 2002; Rogers and Keevil, 1992; Simões et al., 2010). In addition, the composition of the microbial communities in water systems is different in the biofilm and in the water phase (Di Gregorio et al., 2017; Wang et al., 2014). Therefore, analysing the microbial interaction between *Lp* and the resident microbiota in the water and in the biofilm is warranted to fully understand its life cycle and propose better strategies to control its growth.

Pilot-scale water systems have been developed to study disinfection methods (Farhat et al., 2012; Liu et al., 2011; Zhang et al., 2016), *Lp* growth and integration in biofilm (Taylor et al., 2013; Turetgen and Cotuk, 2007), corrosion, scaling, and biofouling (Chien et al., 2012). Of note, *Lp* can be detected in the biofilm in such pilot systems. Nevertheless, few studies have been conducted on pilot cooling towers and, to our knowledge, none accurately depict the complexity of real cooling towers.

Cooling towers are heat exchange devices in which hot water that comes from an external process such as refrigeration, is cooled due to heat exchange between water and air. Hot water is sprayed from the top of the cooling tower by a distribution system through a filling material that breaks the water into small droplets to increase the heat exchange between the air and the water. While water is sprayed, atmospheric air flows from the bottom to the top of the tower. A heat exchange will take place between the air and the water. The water will be cooled and collected at the bottom of the tower and returned to the process that needs cooling. Therefore, a cooling tower system consists of two sections characterized by different temperatures. In addition, the massive input of air in the system increase oxygen availability in the water. It is conceivable that the oxygen concentration is high initially in the basin, but decreases thereafter due to microbial consumption, reaching minimal concentraion at the end of the warm pipe section. As a result, the microbial composition in the biofilm formed on the surface of the different parts of a cooling tower is likely different.

To better understand the growth of *Lp* in cooling towers and its interaction with the resident microbiome, it is therefore crucial to study the biofilm. It is difficult to perform such study on real cooling towers since the pipes are not easily accessible and sampling the biofilm of the pipes requires dismantling the system. As an alternative, we built a lab-scale cooling tower pilot to study the dynamics of *Lp* colonization in relationship to resident microbiota and spatial distribution. This pilot consists of cold and warm water pipe sections connected to an aerated cooling vessel, simulating a typical open evaporative cooling tower. It was filled with water from an operating cooling tower. The objective of the study was to characterize the microbial community in the pilot cooling tower residing in the biofilm and in the water and relate it with the dynamics of *Lp* colonization and local temperature.

## 2. Materials and Methods

### 2.1 Cooling tower pilot

A lab-scale cooling tower pilot was designed to mimic critical components of a real cooling tower (Figure 1). The pilot was installed in a biological safety cabinet to ensure the safety of the laboratory personnel. The system consisted of two symmetrical arrangements of PVC pipes coupled, on one side, to an aerated cooling bioreactor set at 15 °C (Sartorius Stedim Biostat Q Plus, Germany). On the other side, a loop heated by a warm water bath set at 34.4 °C was connected. Each arrangement consisted of eight PVC threaded pipes (McMaster-Carr, USA) of a length of 6 inches and a diameter of 0.5 inch, connected to each other by a threaded T connector and an elbow at the end of each pipe section. A treaded thermocouple type K probe (McMaster-Carr, USA) was fitted in the T connector after the fourth pipe in each arrangement of pipes. The temperature was recorded with a 4-channel portable thermometer/datalogger (OMEGA, USA). A total water volume of 1.05 L was circulated through the pipes by a peristaltic pump using BTP PharMed tubes (Cole Parmer, USA) at a flowrate of 1 L/hr. The temperatures of the water in the pipes were constant during the whole experiment: 22.7 °C for the cool section and 30.7 °C for the warm section. Ambient air was injected in the system at a flowrate of 3 L/min using an aquarium air pump equipped with a 0.2 µm air filter (Millipore, USA). Prior to the start of the experiment, the whole system was disinfected by circulating a 3-ppm sodium hypochlorite solution that was changed every two days. Chlorine residual was measured before changing the chlorine solution using the N,N-diethyl-p-phenylenediamine Colorimetric method 4500-Cl (American Public Health Association (APHA) et al., 2017) and a DR/2010 spectrophotometer (HACH Company, Loveland, CO, USA). In total, two weeks were required to reach stable chlorine residual in the system. Following system disinfection, the pilot was rinsed with unchlorinated sterile distilled water for 24 hours. At this point, the HPC count was 1.4 × 10^3^ CFU/L. The pilot was then filled with water from an actual cooling tower harboring undetectable levels of *Lp* at the time it was collected. An aliquot of water from this cooling tower was kept in a 10 L polypropylene carboy (Nalgene, USA) at room temperature for three months as control water, to distinguish the impact of stagnation from the impact of the pilot system on the water microbiome. After 64 days, the pilot was seeded with *Vermamoeba vermiformis* to a final concentration of 6 x 10^6^ cells/L. At day 72, the pilot was seeded with *Lp* to a final concentration of 3.5 x 10^5^ cells/L. The pilot was dismantled after 92 days, three weeks after the inoculation with *Lp*.

**Figure 1:**
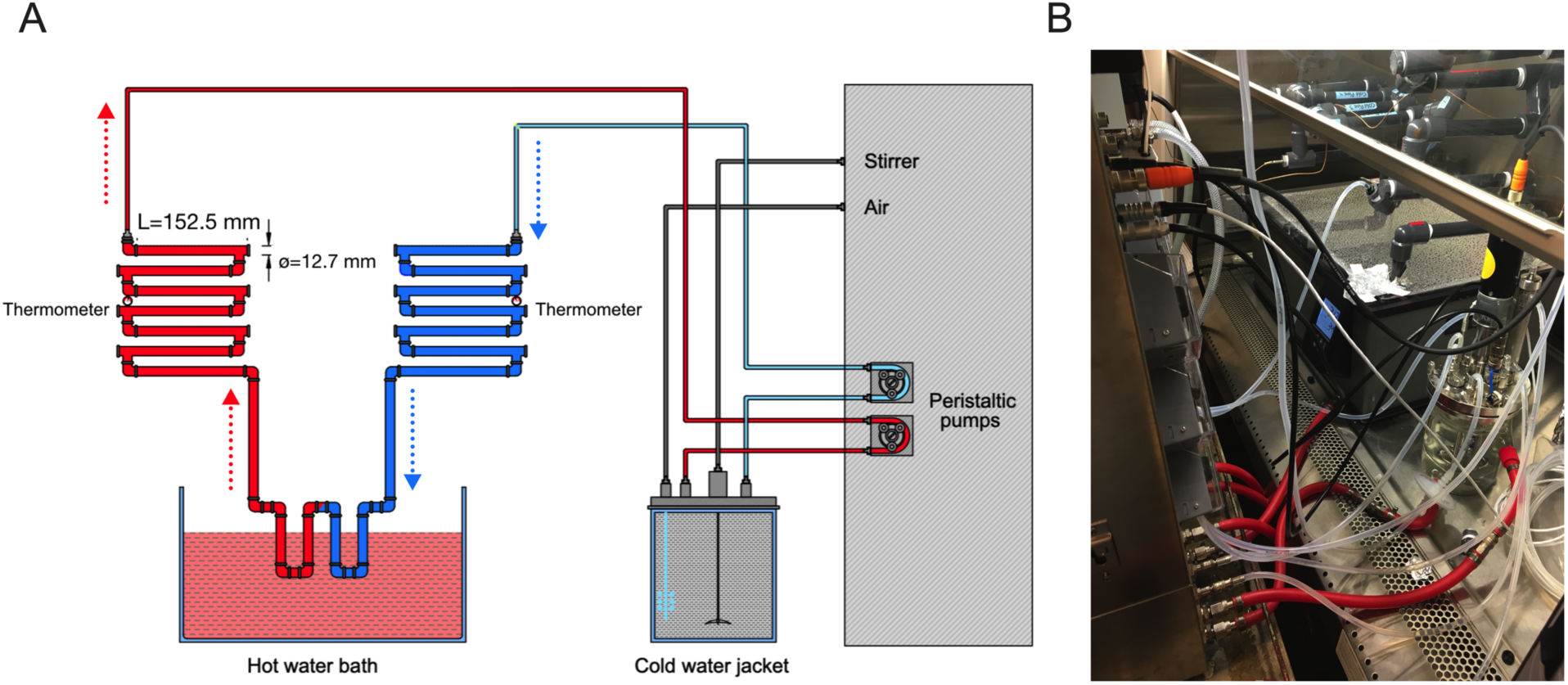
Schematic representation (A) and picture (B) of the pilot tower used in this study. The pilot is composed of a water-jacketed bioreactor vessel connected to a series of cold water pipes (blue), a loop heated by a warm water bath and a series of warm water pipes (red). Water is pumped to the network of pipe and returned to the bioreactor using 2 peristaltic pumps. The bioreactor was maintained at 15°C while the water bath was set at 34.4 °C. The direction of water is indicated with doted arrows.

### 2.2 Inoculation with *V. vermiformis*

*V. vermiformis* (ATCC 50237) was freshly purchased from the American Type Culture Collection and grown in modified PYNFH medium at 30 °C in 75 cm^2^ cell culture flask (Fields et al., 1990). Cells were passaged at a ratio of 1 in 5 when confluence was reached. For inoculation in the pilot cooling tower, the cells were harvested by centrifugation at 800 *g* and washed three times in Page’s Amoeba Saline. The cells were counted with a hematocytometer and a volume corresponding to 6 x 10^6^ cells was added to the pilot through a sampling port in the bioreactor vessel on day 64.

### 2.3 Inoculation with *L. pneumophila*

*Legionella pneumophila* isolated during the Quebec City outbreak in 2012 (*lp120292*) was inoculated in the pilot (Levesque et al., 2014). The strain was maintained at −80 °C in 10% glycerol and grown on BCYE (ACES-Buffered charcoal yeast extract) agar supplemented with 0.25 mg/L L-cysteine and 0.4 mg/L ferric pyrophosphate for 3 days at 37 °C. Several colonies were suspended in filtered sterilized water from the cooling tower to a concentration of 3.5 x 10^5^ cells/mL. One milliliter was added to the pilot through a sampling port of the bioreactor vessel on day 72.

### 2.4 Periodic water sampling

Water sampling was carried out from the bioreactor sampling port. One milliliter samples were taken twice a week for heterotrophic plate count (HPC) on R2A agar. The plates were incubated at 30 °C for 48 hrs. During the first 43 days, a 20 mL sample was collected weekly for DNA extraction. Starting from day 43, the volume was increased to 60 mL. Additional samples of 60 mL were taken after inoculation with *V. vermiformis* and *Lp*. The volume loss was compensated by adding filter sterilized water from the cooling tower that was kept at 4 °C. Due to a considerable decrease in HPC, the volume of sampling was reduced back to 20 mL on day 64 until the end of the experiment. All water samples collected for DNA extraction were filtered through a 0.45 µm pore size filter (Millipore, USA), and the filters were kept at −20°C until DNA extraction.

### 2.5 Pilot disassembly and biofilm sampling

After 92 days of operation, the pilot was disassembled. The water from the bioreactor was first collected. Then, the water was drained from the pipes. The pipes were disassembled, and the attached biofilm was collected as previously described (Proctor et al., 2016; Proctor et al., 2018). Briefly, ten 6-inch pipes (five pipes from the cold part and five from the warm part of the system) were unthreaded. Pipes were capped and filled with 10 mL of 3 mm sterile glass beads. The remaining volume was filled with filter-sterilized water collected from the pilot. The pipes were sonicated for 5 minutes in a sonication bath (Cole Parmer, Canada). Supernatant was collected, and the process was repeated 5 times. An aliquot of the resulting slurry was kept for CFU counts on R2A agar while the rest was filtered through 0.45 µm nitrocellulose filters and kept at −20 °C until DNA extraction.

### 2.6 *16S rRNA* gene amplicon library preparation

DNA was extracted from filters using DNeasy PowerWater Kit from Qiagen (Qiagen, USA), following the manufacturer’s protocol. Each replicate was treated separately. *16S rRNA* gene amplicon sequencing was performed using the dual-index paired-ends approach described by Kozich et al. (2013). Selected samples were analyzed in triplicate. Due to the sampling methodology on the day of dismantlement, the sample of September 13 (day 92) contained detached flocs. Briefly, the extracted DNA was amplified with the 515F and 806R primers targeting the V4 region of the bacterial 16S rRNA gene (Kozich et al., 2013). The PCR amplification was carried out using the Paq5000 Hotstart PCR Master Mix following the manufacturer’s protocol (Agilent, USA). Cycling was performed on an Applied Biosystems Thermal Cycler with cycles consisting of an initial denaturation step at 95°C for 2 min, 25 cycles of 95°C for 2 secs, 55 °C for 15 sec and 72°C for 5 min followed by a final elongation at 72 °C for 10 min. PCR products were purified with AMPure XP beads (Beckman Coulter, USA) according to the manufacturer’s instruction. The purified DNA was quantified with Picogreen using the Quant-iT PicoGreen dsDNA assay kit (Invitrogen, USA). Normalized samples (1.5 ng/µl) were pooled together and mixed with 10% PhiX sequencing control (Illumina, USA). The DNA was diluted to a concentration of 4 pM and denatured with 0.2 N NaOH. The library was sequenced on the MiSeq platform with the MiSeq Reagent V2 250 cycles kit, according to the manufacturer’s instructions.

### 2.7 *18S rRNA* genes amplicon library preparation

*18S rRNA* gene amplicon sequencing was performed using a two-step PCR strategy. Selected samples were analyzed in triplicate. The V9 region of the *18S rRNA* was amplified in a first PCR with primers described in the Earth Microbiome Project protocol (http://www.earthmicrobiome.org/emp-standard-protocols/18s/) (Amaral-Zettler et al., 2009; Stoeck et al., 2010). The cycle of the amplicon PCR consisted of an initial denaturation step at 95°C for 3 min, 25 cycles of 95°C for 30 secs, 55 °C for 30 secs and 72°C for 30 secs, followed by a final elongation at 72 °C for 5 min. PCR products were purified using AMPure XP beads (Beckman Coulter, USA) according to the manufacturer’s instructions. An indexing PCR was next carried out using the Nextera XT Index Kit (Illumina, USA). The index PCR cycle consisted of an initial denaturation step at 95°C for 3 minutes, 8 cycles of 95°C for 30 secs, 55 °C for 30 secs and 72°C for 30 secs, followed by a final elongation at 72 °C for 5 min. Both PCR amplifications were carried out using the Paq5000 Hotstart PCR Master Mix (Agilent, USA). The library was sequenced on the MiSeq platform with the MiSeq Reagent V2 250 cycles kit, as described above for *16S rRNA* gene amplicon sequencing.

### 2.8 Data Processing

The raw sequence reads were deposited in Sequence Read Archive under the BioProject accession number PRJNA588467. The sequenced reads were processed using the Mothur pipeline (Schloss et al., 2009). Paired-end reads were first assembled into contigs. Contigs that presented ambiguous bases or that were longer than 275bp for the *16S rRNA* gene sequencing and 373 bp for the *18S rRNA* gene sequencing were removed. The SILVA 132 database was used to align the sequences. Ends and gaps were trimmed in order to have the same alignment coordinates for all the sequences. Chimeras were removed using the VSEARCH algorithm. Two of the replicates for the *16S rRNA* gene analysis had significantly lower read counts than the rest; hence, they were removed from the analysis (Warm pipe 7 c, Cold pipe 7 b). The rest of the samples were rarified to the next sample with the lower number of reads (3038 read counts). For the 18S *rRNA* gene analysis, the samples included in the analysis were rarified to the lowest read count sample (6325 read counts). For *16S rRNA* gene sequencing, non-bacterial sequences such as Eukaryotes, chloroplasts, Archaea and Mitochondria were removed. For *18S rRNA* gene sequencing, only Eukaryotic sequences were considered. For *18S rRNA* gene sequencing, only Eukaryotic sequences were considered. Operational Taxonomic Units were defined at an identity cut-off of 97%, by assigning the OTUs de novo. The OTU data was analyzed with the MicrobiomeAnalyst web-based tool (Dhariwal et al., 2017) . Default parameters were used to filter OTU with low counts (OTUs with less than 2 counts in at least 20% of the samples were removed). Beta diversity was calculated with the Bray-Curtis dissimilarity index to analyze differences between samples. Non-metric multidimensional scaling (NMDS) and principle coordinate analysis (PCOA) were used to visualize the data. PERMANOVA analysis was performed to analyze the statistical significance between groups.

## 3. Results and Discussion

### 3.1 Physical and microbial characteristics of the pilot

The pilot was designed to mimic as accurately as possible the operation of a real cooling tower. The temperatures in the cold and hot pipe were remarkably stable at 22.7 °C and 30.7°C respectively, reproducing the temperature typically seen in a cooling tower (ASHRAE, 2008). The pH was also stable around 8.1 during the whole experiment. The pilot was seeded with water collected from an operating cooling tower and filter-sterilized water from that tower was used as make up water.

HPCs in the reactor water were between 10^5^ and 10^6^ CFU/mL during the first forty days of the experiment, showing a relative stability (Figure 2A). A decrease in the HPC was noticed between day 43 and 70. During this period, the volume of water collected from the bioreactor for DNA extraction was increased from 20 mL per week to 60 mL, which increased the addition of makeup water. This apparently caused over dilution of the microbial population in water. A rise of the CFU in the water was observed when the volume taken was decreased back to 20 mL around the time of inoculation with *V. vermiformis* on day 64. Inoculation of *Lp* did not seem to affect CFU counts. On the last day (92), there were 1.40 × 10^5^ CFU/mL in the water, for an estimated total cultivable biomass of 1.47 × 10^8^ CFU in the system, assuming a volume of water of 1.05 L. Of note, some flocs were visible in the water collected from the bioreactor on the last day.

**Figure 2:**
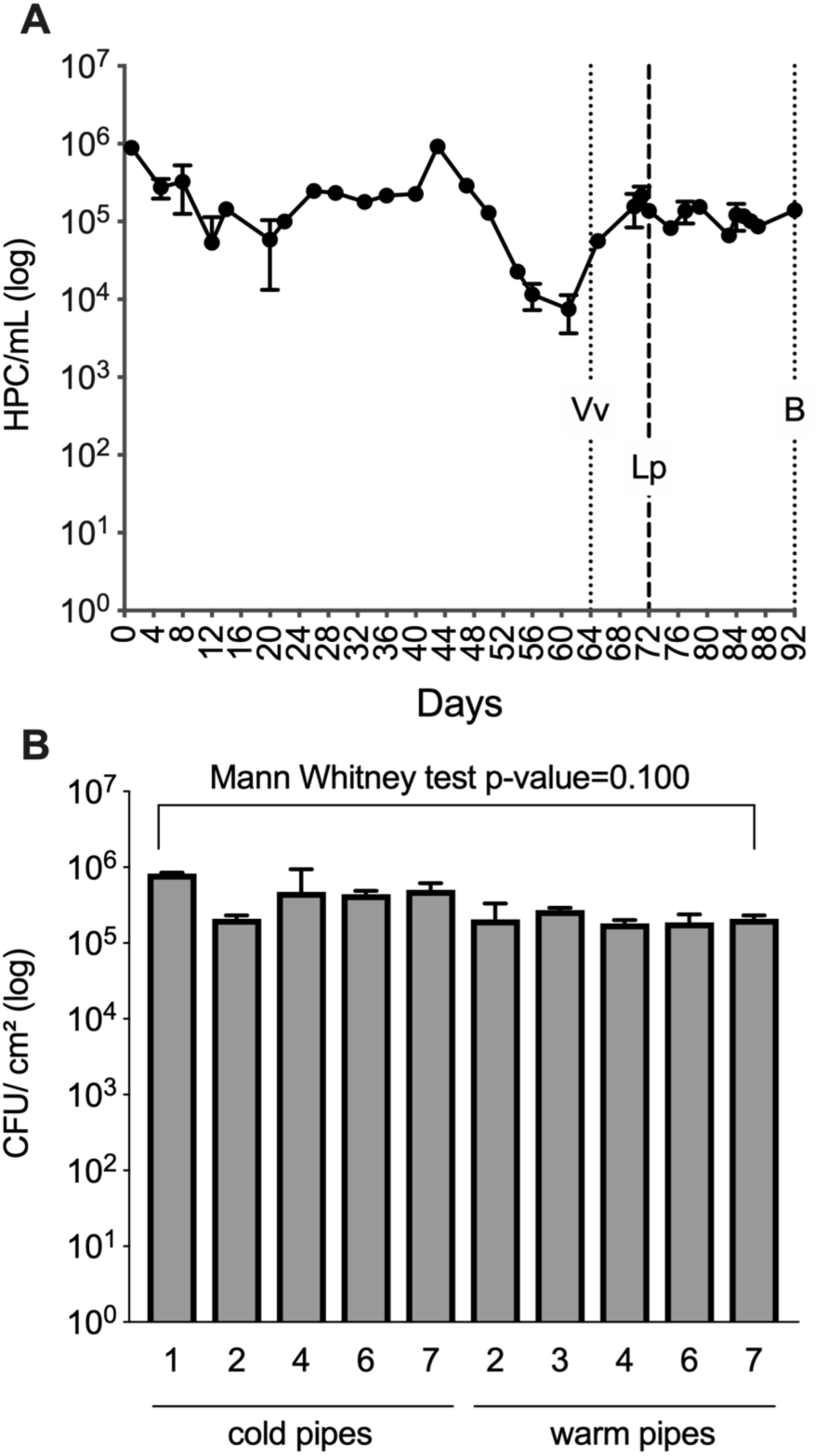
The system was monitored by heterotrophic plate count during operation and at the time of dismantlement. Water samples of 1mL were taken twice a week for 92 days and HPC counts were performed on R2A agar (A). Data represents the mean of triplicate samples with standard deviation. The time of inoculation with *V. vermiformis* (Vv) and with *Lp* (Lp) is indicated. On day 92, the system was dismantled and biofilm (B) samples from five pipe segments from the cold section and from the warm section were harvested by sonication with glass beads and HPC was determined on R2A agar (B). Data represents the average of CFU per cm² with standard deviation. Statistical significance between biofilm grown at the different temperatures was determined using a Mann-Whitney test.

Biofilms were extracted from the pipes using a sonication and glass beads method (Proctor et al., 2016; Proctor et al., 2018). There was no significant difference (Mann-Whitney test) between the HPC counts from the biofilm samples taken from the cold pipes and the ones taken from warm pipes (Figure 2B). The average cultivable biomass in the biofilm was 3.51 × 10^5^ CFU/cm^2^. Using an estimated surface of 2257.5 cm^2^ for the pipe system, the total cultivable biomass present in the biofilm is estimated to 7.92 × 10^8^ CFU. This is at the upper range of what was previously reported for biofilm sampled inside drinking water distribution systems (Wingender and Flemming, 2011). There were only 5 times more cultivable microorganisms in the biofilm than in the water of our pilot system at the time biofilm was sampled (day 92). This is not consistent with the literature reporting that about 95% of bacterial cells in water systems are fixed on surfaces (Flemming et al., 2002). This can be due to the fact that the temperature range in the system and the lack of disinfection methods was ideal for the organisms to be in the planktonic state. Alternatively, the relatively high surface volume ratio of our system (2.15 cm^−1^) is known to promote cell release from the biofilm into the water (Bedard et al., 2018)

### 3.2 Characterization of the eukaryotic and bacterial communities in the pilot cooling tower

Bacterial community profiling was carried out on the samples collected on day 1, 57, 79, 83, 86 and 92. The bacterial community of the water changed drastically between day 1 and day 57 but seemed relatively stable afterward. The inoculation with *V. vermiformis* and *Lp* had only a minor effect on the general composition of the water microbiota (Figure 3A). This indicate that the initial inoculated microbiota was modified by the system during the first two months and eventually reached a relative equilibrium by day 57. *Obscuribacterales* and *Verrucomicobiceae* were the most predominant bacterial families in the water samples, with relative abundance going from 7.7% to 27.9% and 8.8 to 22.2% respectively, excluding the samples taken at inoculation. Several water samples analyzed with 18S rRNA gene amplicon sequencing produced very low number of reads and were therefore excluded from the analysis. *Poterioochromonas* was the most predominant eukaryotic genus in the water samples taken on day 57 but was reduced to 15% on day 92, apparently being replaced by other organisms. Interestingly, *Poterioochromonas* is a flagellated protist that preys on other microbes, including bacteria (Saleem et al., 2013). Possibly, its favorite prey type disappeared from the system during day 87 and day 92, possibly because of over predation, which resulted in a decline in the population of *Poterioochromonas*. Alternatively, it might be a previously unknown host of *Lp*, since the decline of *Poteriochromonas* coincide with the growth of *Lp*. Additional experiments will be required to confirm this hypothesis. *Vermamoeba* was already present at day 57, before inoculation with *V. vermiformis*, and was also present at day 92. Other OTUs harboring potential hosts for *Lp* were also detected such as *Oligohymenophorea* and *Naegleria*, but only at the later time point (Figure 3B). In the biofilm sampled on day 92 (Figure 3C), *Nitrosomonadaceae* was the most predominant bacterial family in the warm pipes (17.1% to 22.5%). In contrast, the levels of *Burkholderiaceae* (9.3% to 22.2%), *Microscillaceae* (7.9% to 16.8%) and *Rhodocyclaceae* (8.0% to 17.1%) seemed higher in the biofilm formed in the cold pipes. *Oligohymenophorea* was the most abundant eukaryotic genus in the biofilm samples (Figure 3D), having a higher abundance in the biofilm formed in the cold pipes (31.3% to 73.7%) compared to the hot pipes (3.1% to 18.5%). *Vermamoeba* and *Naegleria* were also detected in the biofilm samples in pipes at both temperatures.

**Figure 3:**
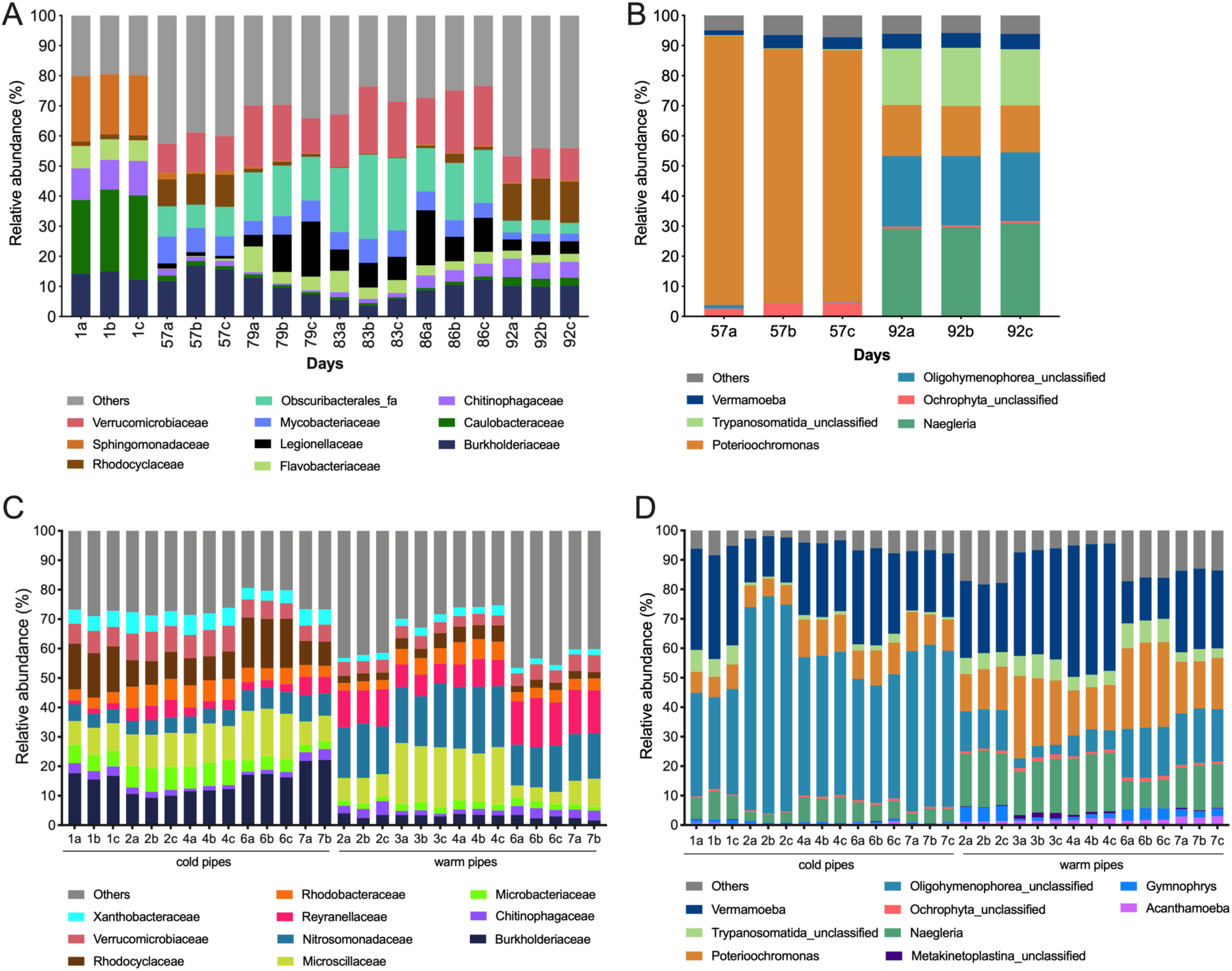
The microbial community composition of water samples (A and B) and biofilm samples (C and D) was determined. The bacterial community was analyzed by *16S rRNA* gene amplicon sequencing (A and C) while the eukaryotic community was determined using *18S rRNA* gene amplicon sequencing (B and D). The data are presented as the relative abundance of OTUs classified at the most appropriate taxonomic level.

It is difficult to compare the microbiome of this pilot cooling tower with other studies since water source and regional climate shape the microbiome of cooling towers (Llewellyn et al., 2017; Paranjape et al., 2020). Nevertheless, some similarities are observed between our pilot cooling tower and other studies. For instance, our results showed that *Burkholdericeae* was abundantly found in the water of the pilot cooling tower. This is in agreement with other studies reporting the microbial communities of similar environments (Paranjape et al., 2020; Tsao et al., 2019). *Verrumicrobiaceae* was also an abundant family in the system, which is consistent with its presence in natural water reservoirs (Boucher et al., 2006; Zwart et al., 2002). Taxa previously identified as organisms capable of forming biofilm such as *Pirellulaceae, Rhodobacteraceae and Caulobacteraceae,* were identified in the biofilm samples of the pilot (Elifantz et al., 2013; Entcheva-Dimitrov and Spormann, 2004; Miao et al., 2019).

Beta diversity analysis was used to evaluate the effect of the pilot on the microbiota by comparing the bacterial communities of the pilot tower water at day 92, of the initial water used for inoculation, and of the initial water incubated at room temperature on the bench for 92 days (Figure 4). The microbiome of the pilot water was significantly different than the initial water used to seed the system and shows a significant difference from the microbiome of the stagnant water (PERMANOVA F-value: 269.21; R^2^: 0.98901; p-value < 0.001). This result indicates that specific characteristics and operating parameters of our pilot tower, such as temperature, dissolved oxygen, and water flow, shaped the resident microbiota. These parameters were identified as the main factors influencing the resident microbiota of a model water distribution system (Douterelo et al., 2017).

**Figure 4:**
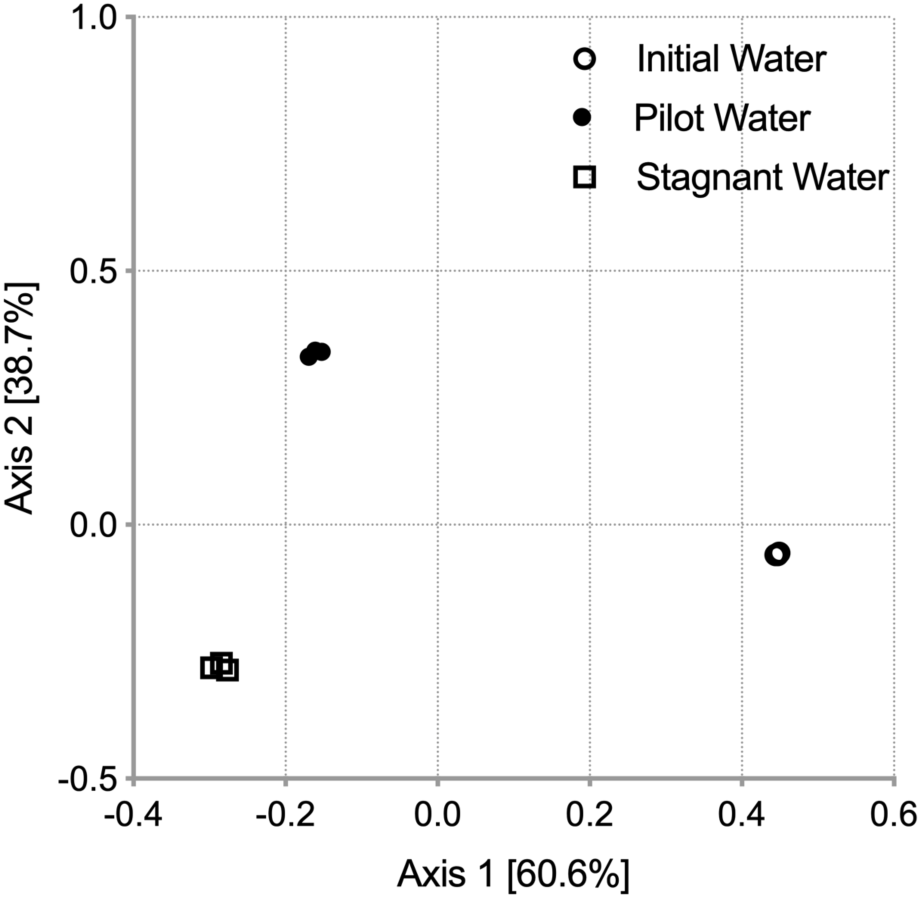
Beta-diversity was used to analyze the effect of the pilot cooling tower on the bacterial community. A principal coordinate analysis (PCoA) plot of bacterial profiles of the water samples from the cooling tower at day 92 (pilot water), from the initial water (day 1) and control stagnant water was used. Statistical significance was determined using PERMANOVA.

### 3.3 Presence of *Legionella* in the system

The presence of *Legionella* in the system was evaluated using the results of the *16S rRNA* gene amplicon sequencing (Figure 5). The relative abundance of *Legionella* in the water at the beginning of the experiment (day 1) was almost null (0.02%). An increase in the relative abundance of *Legionella* was observed after 57 days reaching 3.0%. Right after the inoculation with *Lp* on day 72, the relative abundance of *Legionella* in water was 11.0 %. reaching 13 % at day 86, but then dropping to 4% on day 92 (Figure 5B). The relative abundance of *Legionella* in the biofilm samples was extremely low, but detectable (Figure 5B). One of the objectives of this study was to observe the spatial distribution of *Legionella* within cooling towers. While we were expecting to see significant differences in the relative abundance of *Legionella* in biofilm at different temperatures, this was not observed. It is tempting to conclude that most of *Legionella* was in the water phase in the system. The presence of *Legionella* in biofilms within water distribution systems has been reported in several studies (Abdel-Nour et al., 2013; Abu Kweek and Amer, 2018; Armon et al., 1997; Buse et al., 2014; Buse et al., 2012; Declerck, 2010; Lau and Ashbolt, 2009; Moritz et al., 2010; van der Kooij et al., 2005). Several factors influence the formation of biofilm by *Legionella* and its ability to integrate biofilms (Buse et al., 2017; Piao et al., 2006; Rhoads et al., 2017; Rogers et al., 1994a). Indeed, *Legionella* incorporates in pre-established biofilms as a secondary colonizer. Instead of attaching to surfaces and growing biofilm, the bacterium will form an association with other microbes that previously developed biofilm (Buse et al., 2017) . Thus, integration of *Legionella* into biofilms is affected by water temperature, surface material, water quality, microbial composition of the biofilm and biofilm age (Buse et al., 2017).

**Figure 5:**
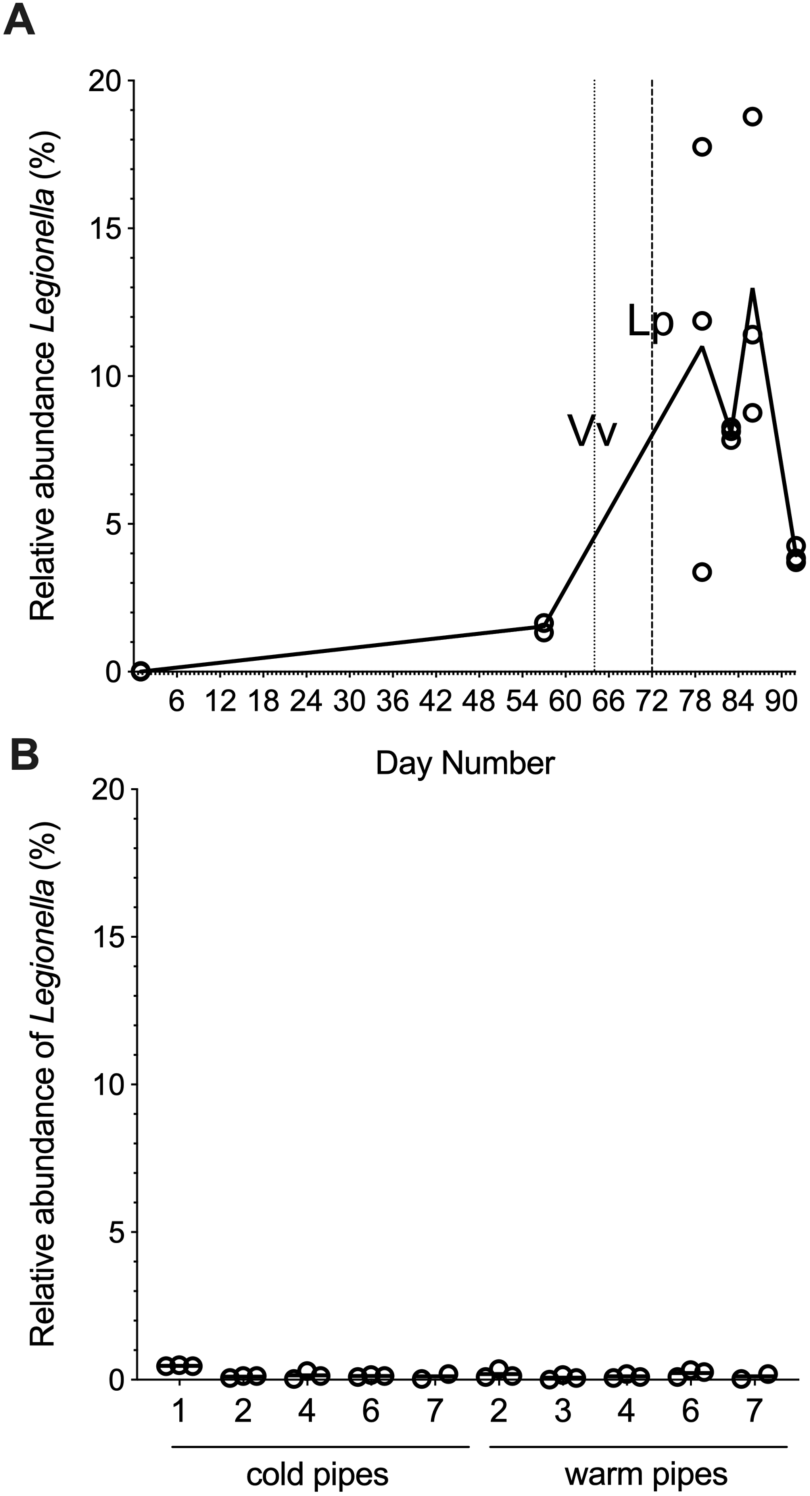
The relative abundance of *Legionella* in the pilot water (A) and biofilm (B) was determined from the *16S rRNA* gene amplicon sequencing data. The percentage of abundance of the reads of the *Legionella* OTU was calculated according to the rarefied number of reads for each sample after rarefaction. Individual replicates are shown. The line in (A) connects the means of each time points.

Potential host of *Legionella*, such as *Vermamoeba*, *Acanthamoeba*, *Naegleria* and ciliates (*Oligohymenophorea*) were detected in the water samples (Figure 3B) as well as in the biofilm. Intracellular growth of *Lp* in biofilm is dependent on the concentration of host cells (Shaheen et al., 2019). The presence of host cells in the biofilm as well as the temperature being between 22.7 and 30.7°C in the pipes (Figure 3B) suggest that *Legionella* had ideal growth conditions (Ashbolt, 2015; Fields et al., 2002; Moffat and Tompkins, 1992; Rowbotham, 1980).

Furthermore, it was previously shown that *Legionella* is able to integrate biofilm formed on PVC, the material used for the pipe in our studies (Armon et al., 1997; Rogers et al., 1994a). Therefore, we were expecting to find a larger proportion of *Legionella* in the biofilm than in the water. However, the conditions found in the water of our pilot, including the presence of specific host species, might be ideal for *Lp* growth, favoring its presence in water. This is supported by the presence of host cells in the water phase. The lack of time points for the analysis of the composition of the biofilm prevents us from making any assumptions about the dynamics of the microbiota in the biofilm. It is possible that *Legionella* concentration in the biofilm was higher at an earlier time point. The spatial distribution of *Lp* in cooling towers will need to be studied further.

### 3.4 Difference in the water and biofilm communities

At first sight, the composition of the microbial communities of the water and of the biofilm seems different. To characterize the communities further, the Shannon Index was calculated to measure alpha diversity (Figure 6A and B) while beta diversity was used to assess dissimilarities between the communities (Figure 6C and D). For the water samples, the analysis was performed only with the samples from day 79 to 92, to avoid changes induced by inoculations of *V. vermiformis* and *Lp*. There was a slight but significant difference between the alpha diversity in the biofilm samples and in the water samples for the bacterial communities but not for eukaryotic communities. Beta diversity analysis revealed that the water and biofilm bacterial communities were dissimilar, clustering in distinct groups (Figure 6C, PERMANOVA F-value: 36.174; R^2^: 0.488; *P* < 0.001; Stress = 0.0765). This was also observed for the eukaryotic communities Figure 6D, PERMANOVA F-value: 11.076; R^2^: 0.246; *P*< 0.001; Stress = 0.0945; however, the samples from day 92 clustered with the biofilm samples (open circle samples in Figure 6D). This could be due to contamination of the water samples with biofilm fragments as flocs were observed in the bioreactor during dismantlement. Similarly, the bacterial communities at day 92 seems to share characteristics between the water and biofilm group (Figure 6 C), although the similarity was less pronounced then for the eukaryotic community.

**Figure 6:**
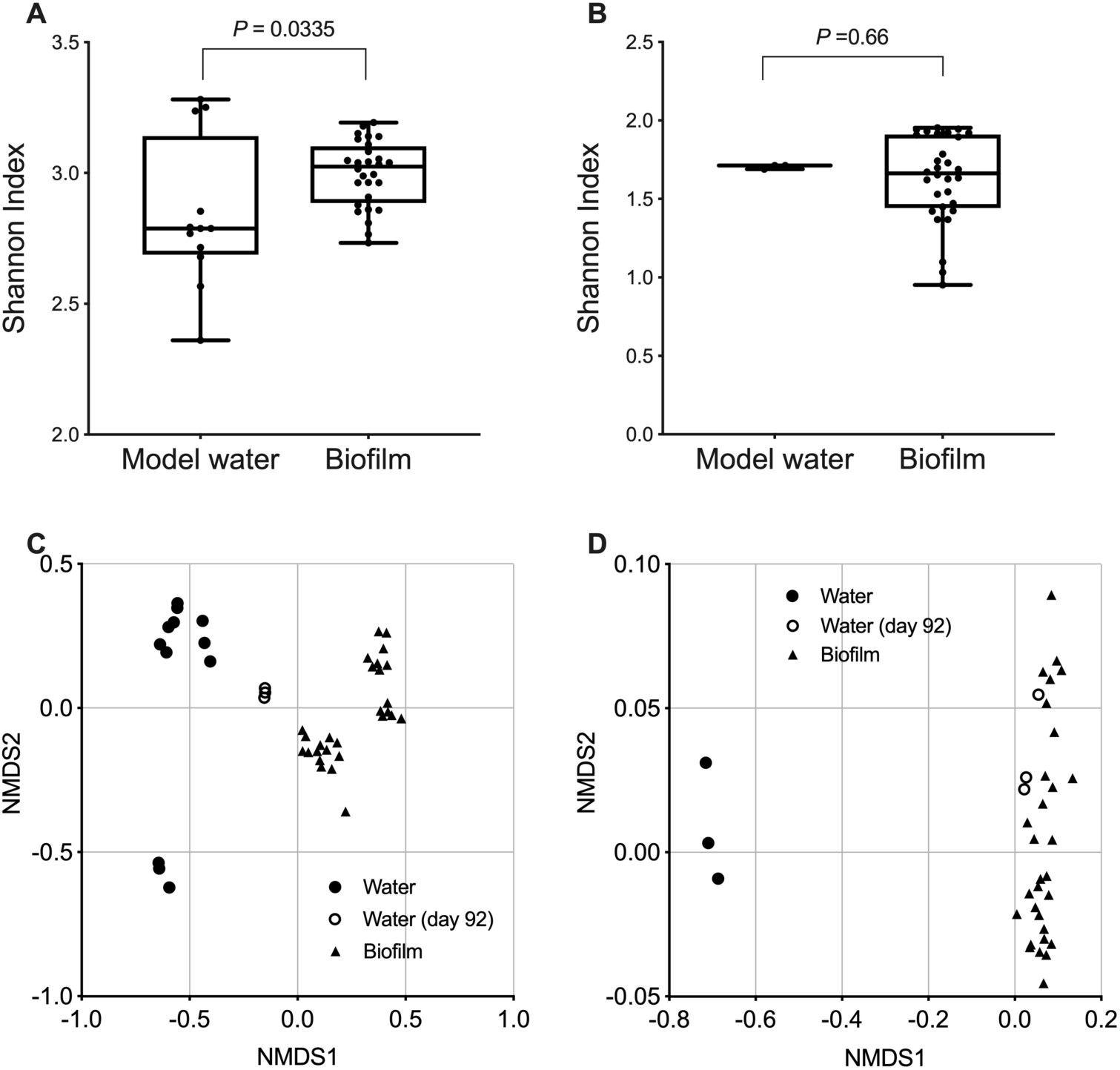
Alpha diversity of samples from the pilot for the bacterial (A) and eukaryotic community (B), categorized by the type of samples: water and biofilm. A Mann-Whitney test was performed to determine the statistical significance. Beta diversity was calculated for the bacterial (C) for the eukaryotic (D) community categorized by the type of samples. PERMANOVA was used to assess statistical significance. Only day 79 to day 92 were analyzed for the bacterial community of the water to avoid noise introduced by addition of *V. vermamoeba* and *Lp*.

Next, the machine-learning algorithm LEfSe was used to identify bacterial and eukaryotic taxa associated with either the water samples or the biofilm samples (Segata et al., 2011). Only water samples taken after day 79 were considered for the LEfSe analysis of bacterial communities. The algorithm was able to identify significant taxa associated with water and biofilm (Figure 7). Of note, *Legionellaceae* were enriched in the water while its hosts, including *Vermamoeba*, *Acanthamoeba,* and *Oligohymenophorea*, were enriched in the biofilm. The enrichment of amoebas in the biofilm of our pilot system is consistent with what was previously reported for the biofilm in water distribution systems (Taravaud et al., 2018; Thomas et al., 2004; Thomas et al., 2008). Therefore, it seems that *Legionella* is mostly in the water while its hosts are mostly in the biofilm, which seems counterintuitive. A possible explanation is that *Legionella* actively grows in the biofilm, where the hosts are located, but it is released into the water after intracellular replication, as previously shown (Greub and Raoult, 2004; Lau and Ashbolt, 2009). The bacterium can also be expelled in cysts from ciliates and amoeba such as a *Tetrahymena* and *Acanthamoeba*, respectively (Berk et al., 2008; Bouyer et al., 2007; Hojo et al., 2012).

**Figure 7:**
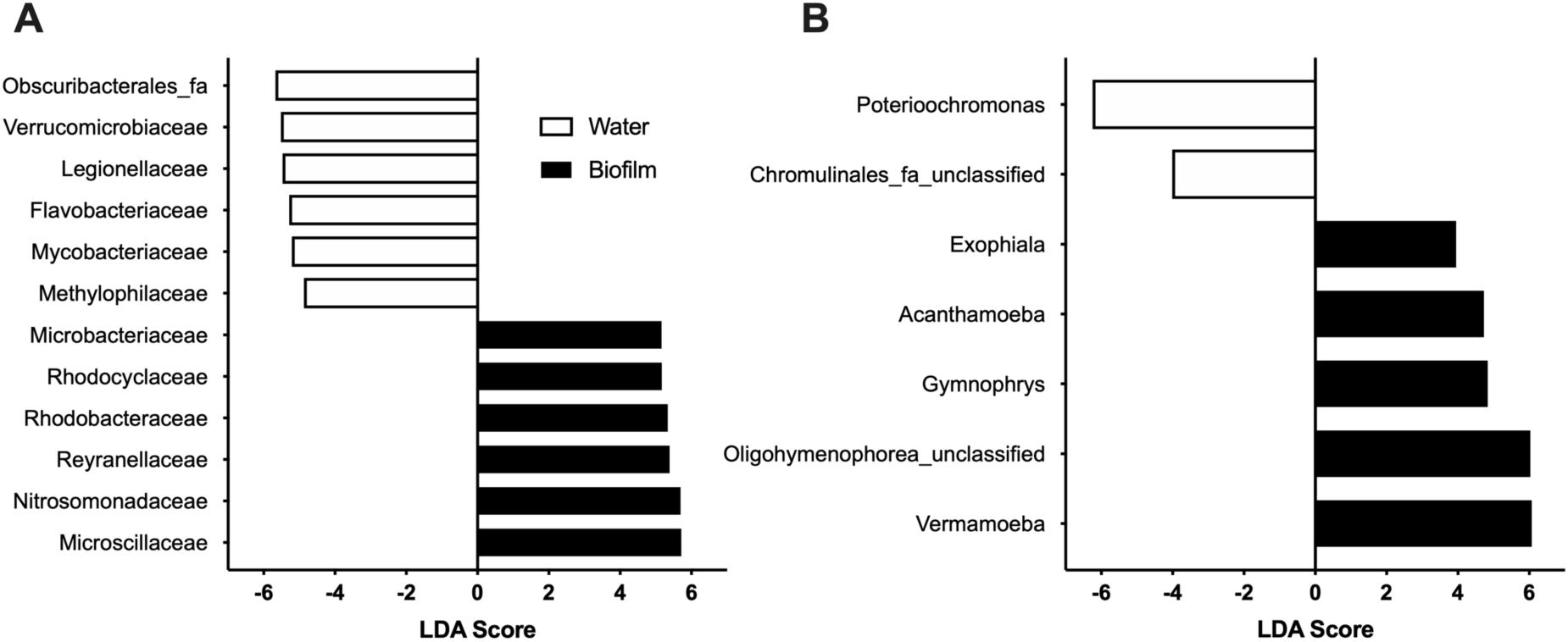
The machine learning algorithm LEfSe was used to identify significant bacterial (A) and eukaryotic (B) taxa associated with either water (open bars) or biofilm (black bars). Only significant taxa (*P* < 0.02) are shown.

### 3.5 Influence of the temperature on bacterial and eukaryotic communities in the biofilm

Beta diversity was calculated to analyze the difference between biofilm samples (Figure 8). Grouping the samples according to the temperature produced significantly different clusters for the bacterial (Figure 8A, PERMANOVA F-value: 37.838; R^2^: 0.59272, *P* < 0.001, Stress = 0.10321) and for the eukaryotic communities (Figure 8B, PERMANOVA F-value: 37.717, R^2^: 0.57393, *P* < 0.001; Stress = 0.15982). This is not surprising since temperature is known to affect biofilm formation and composition, as well as the presence of *Lp* (Buse et al., 2017). The strength of our study is that our unique pilot design allows us to decipher the effect of temperature in a single system were the different surfaces are inoculated with the same microbiota. The specific biofilm communities present at the different temperatures likely established gradually from the original inoculum eventually reaching a specific composition. It is not clear if the composition of the biofilms was stable at the time of disassembly. A time course study will need to be performed to understand the dynamic of biofilm establishment at different temperature in the same system. To our knowledge, there is a scarcity of study assessing this particular point. Next, LEfSe was used to identify bacterial and eukaryotic taxa enriched in biofilm at 22.7 °C and at 30.7 °C. Bacterial families such as *Burkholderiaceae, Rhodocylaceae* and *Microbacteriaceae* were predictive of biofilm at 22.7 °C while *Nitrosomonadaceae* and *Reynellaceae* were predictive of biofilm at 30°C. Interestingly, the ciliate genus, *Olygohymenophorea* was predictive of cold biofilm while amoeba such as *Naegleria* and *Acanthamoeba* were predictive of warm biofilm. It is possible that the species of *Oligohymenophorea* present in the system have an optimal growth temperature closer to 20°C. Indeed, a recent study of the microeukaryote communities in the St-Charles river in Quebec, Canada, revealed that ciliates are more abundant during the winter period (Cruaud et al., 2019). It is tempting to speculate that ciliates might be more important for intracellular growth of *Lp* at low temperature and amoebas at higher temperature in water systems. The role of ciliates in the life cycle of *Legionella* in water systems running at low temperature should be investigated further.

**Figure 8:**
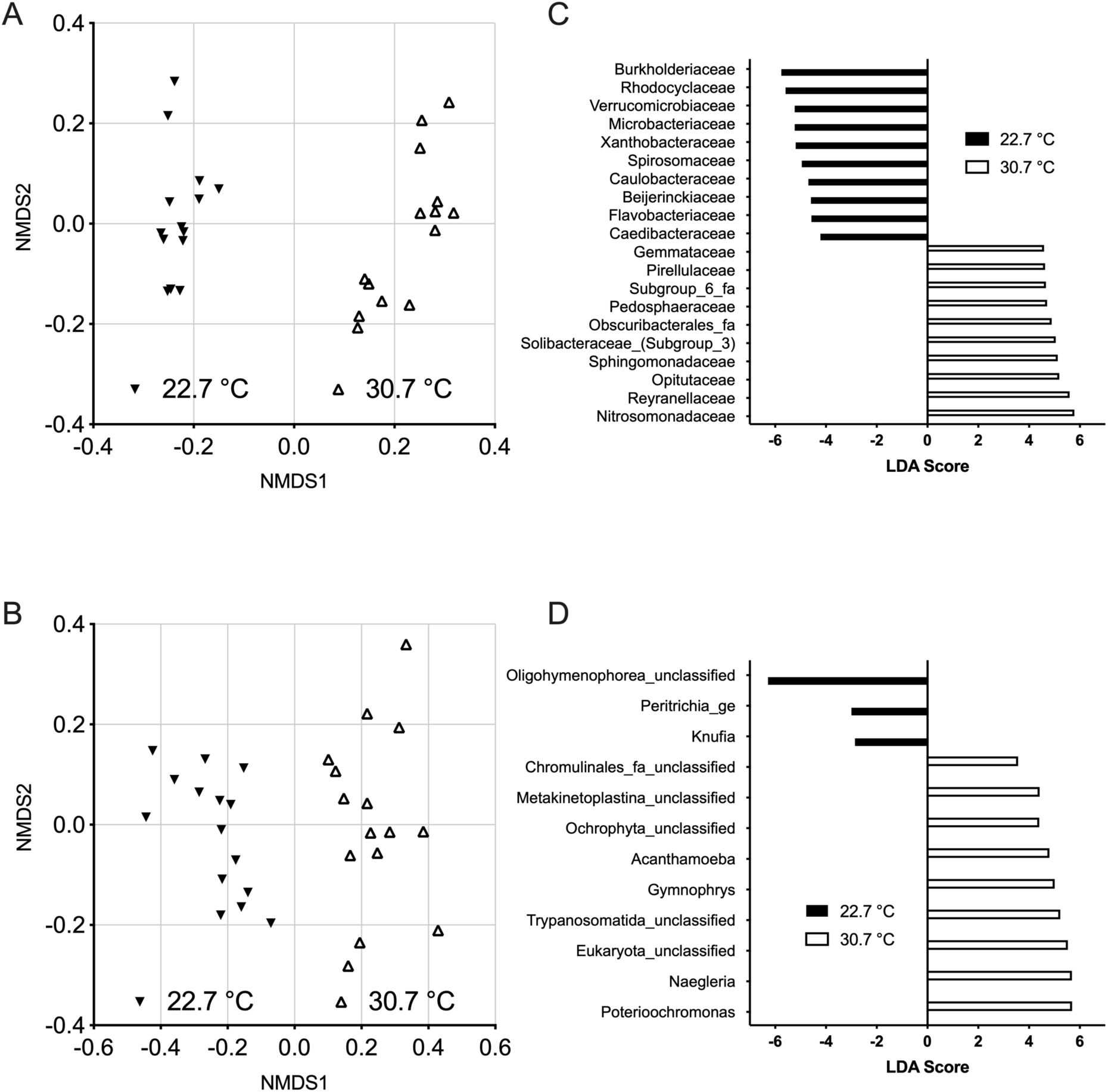
Temperature affects the microbial composition of the biofilm. Beta-diversity of the bacterial (A) and eukaryotic (B) communities was calculated using the Bray-Curtiss index and a non-metric dimensional scaling (NMDS) plot. The samples were grouped by temperature. PERMANOVA was used to assess statistical significance. A LEfSe analysis was performed to identify bacterial (C) and eukaryotic (D) taxa associated with each temperature. Only statistically significant taxa (*P <*0.02) are shown.

## 4. Conclusion

This study illustrates the importance of studying the microbial composition of the water as well as the biofilm to fully understand *Legionella* ecology in water systems. From our study, three main observations emerge.

- In our pilot, the temperature had a great impact in the composition of the resident microbiota of the biofilm, indicating that the cold and warm pipe section of actual cooling towers are likely to harbor different microbial population.
- The host cells were mainly present in the biofilm, while *Legionella* was present in a lower proportion in the biofilm at the time of sampling. This suggest that *Legionella* grows in the biofilm but is released back in the water afterward, following a host-prey cycle within hosts population.
- Ciliates and amoebas seem to inhabit different parts of the system, the former preferring the colder part. Therefore, additional research is needed to appreciate the role of ciliates in *Legionella* growth at lower temperature. Finally, our study supports the usefulness of pilot systems in studying the ecology of *Legionella* and other water-borne pathogens.

## Acknowledgments

We would like to thank Michel Gauthier for his help in sampling the cooling towers used to inoculate our pilot systems and Jose Antonio Torres for his help creating the pilot diagram. This work was supported by a FRQNT Team grant (2016-PR-188813) and a NSERC Discovery Grant (RGPIN/04499-2018) to SPF. ATP was funded by a scholarship from CONACYT. Mengqi Hu was funded by a MITACS Globalink award.

